# Electrical impedance detection of sickle cell vaso-occlusion in microfluidic capillary structures

**DOI:** 10.1101/2020.07.29.227215

**Authors:** Yuhao Qiang, Jia Liu, Darryl Dieujuste, E Du

**Affiliations:** Department of Ocean and Mechanical Engineering, Florida Atlantic University, Boca Raton, FL 33431, USA; Department of Biological Sciences, Florida Atlantic University, Boca Raton, FL 33431, USA

## Abstract

Sickle cell disease (SCD) is primarily associated with episodic vaso-occlusive events. Poorly deformable sickle cells may get stuck in small blood vessels, slow down or block blood flow, leading to local hypoxia that damages tissues and organs. In this paper, we present a novel electrical impedance sensing technique for detection of the progressive occlusion by sickle cells in microfluidic capillary structures. Changes in both resistance and reactance of the sickle blood flow were observed at multiple low frequencies (< 500 kHz), upon the deoxygenation and reoxygenation processes. In contrast, no obvious impedance changes were observed in the flow of normal blood cells and sickle blood cells treated with anti-sickling agent. Accuracy of the impedance-based detection of the vaso-occlusion process was verified by microscopic observation. The results show the distinct sensing performance of sickle cell vaso-occlusion by electrical impedance, which does not require sophisticated optical microscopy or video processing. The low frequency impedance sensing can be achieved by replacing the benchtop equipment with low-cost, high precision impedance converter system, allowing for detection of sickle cell vaso-occlusion in point-of-care settings.

## Introduction

Vaso-occlusion is responsible for a variety of clinical complications in sickle cell disease (SCD) such as pulmonary hypertension, stroke and organ damages. Current evidence indicates that vaso-occlusion is the most conceivable etiology of sudden death such as of competitive athletes with sickle cell trait during periods of extreme physical exertion in sports.^1^ The defective gene responsible for SCD leads abnormal hemoglobin (HbS) in patient’s red blood cells (RBCs), also known as sickle cells, which are stiffer and stickier than normal RBCs. In particular, Hbs in sickle cells can polymerize when exposed to low oxygen concentration, causing misshaped cell membrane and reduced cell deformability. Vaso-occlusion process is known to be multifactorial,^2^ where leukocyte adhesion, vascular intimal hyperplasia and fat embolism act as prerequisites, and trapping of poorly deformable sickle cells may ultimately result in obstruction of small blood vessels and stop the blood flow. Timing is critical for most of sickle cells escaping the narrow capillaries before they become rigid enough to get trapped.^3^ Hence, the delay time in HbS polymerization and cell sickling is likely a key parameter predictive of vascular occlusion.^4^

SCD management requires patient self-monitoring by home diary of pain and stress as suggested by the US Centers for Disease Control and Prevention.^5^ For patients with SCD, who require treatment with some anti-sickling therapies, it is essential that care providers can rapidly monitor their ability to maintain healthy homeostasis while preventing vaso-occlusion. However, these pain measurements are subjective and unquantifiable.^6^ There is a urgent need for convenient tools for monitoring and prediction of the vaso-occlusive pain, as well as patient’s therapeutic outcomes. Existing laboratory techniques for monitoring SCD used in hospital include hemoglobin electrophoresis, isoelectric focusing, and cation-exchange high-performance liquid chromatography, require sophisticated equipment, special reagents and complicated procedures.^7^ Growing efforts have been made in developing affordable point-of-care devices, such as a high-throughput screening assay based on detecting the ability of RBCs to traverse a column of tightly packed Sephacryl chromatography beads,^8^ and a paper-based colorimetric assay which can qualitatively differentiate SCD blood from sickle cell trait and normal blood using the blood stain patterns, as well as sensors for monitoring oxygen saturation levels in blood.^9^ Additionally, the rapid development of microfluidic assays enables the *in vitro* cell sickling measurement from the sub-cellular level to the whole blood level, such as the deformability measurement of cellular membrane,^10^ microfluidic assays for bio-rheological characteristics of sickle RBC suspension^3, 11^ and single RBCs detection of HbS polymerization in deoxygenated liquid drops,^12^ investigation of pathologic adhesion in blood rheology^13^, and image flow cytometry for morphologic characteristics of deoxygenated RBCs.^14^ Unfortunately, these in vitro assays do not offer capability in direct and quantitative measurements of vascular occlusion induced by rigid sickle cells.

To that end, we developed an *in vitro* vaso-occlusion assay with capillary-inspired structures allowing direct observations of biorheology and vascular obstruction of individual SS RBCs under microscope.^4^ To overcome the limitation of the blood testing and the requirements of microscopic observations and video processing, we facilitate the vaso-occlusion assay by integration of an electrical impedance sensor ^15, 16^ in the microfluidic channel. In this paper, we demonstrate the capability of this new assay for a real-time measurement of sickle cell traversing through microcapillaries including the progressive vaso-occlusion upon deoxygenation and resume of blood flow upon reoxygenation. We further evaluate the effects of the anti-sickling drug on the blood flow in the microfluidic assay. We envision that this method can be further developed for use in the point-of-care settings by replacing the benchtop impedance measurement system with low-cost impedance converter as in the portable impedance flow cytometry device.^17^

## Study design

### Sample preparation

Normal blood and sickle cell blood samples were obtained following Institutional Review Board (IRB) approvals from Florida Atlantic University and the University of Miami. All samples were stored at 4°C and tested within one week of collection. Prior to each experiment, blood samples were washed twice with phosphate-buffered saline (PBS) at the speed of 2000 rpm at room temperature for two minutes. The hematocrit of each tested sample was adjusted to be 0.1% by resuspending 1 μL RBC pellet into 1 mL in PBS. For the testing of anti-sickling effect of 5-hydroxymethyl-2-furfural (5-HMF) on the vaso-occlusion, RBC suspensions were incubated with 5-HMF (5 mM) for 60 min at 37 °C in 1.5 ml Eppendorf tubes. The treated cells were washed twice with PBS before the measurement.

### Experimental Setup

Fig. 1A provides a schematic diagram of the overall experimental setup. Gravity-driven flow approach was used to generate the flow of RBC suspension in the cell channel. An equivalent pressure difference (∼500 Pa) was created by connecting the cell channel to the external hydraulic columns via flexible Tygon microbore tubing (0.020 “i.d.× 0.060”o.d.). Transient hypoxia condition in the cell channel was created by gas diffusion through a 150 μm thick polydimethylsiloxane (PDMS) layer at 4.5 psi by switching the gas mixture supplies in the gas channel from high oxygen concentration (17.5% oxygen, 5% carbon dioxide with the balance of nitrogen) to low oxygen concentration (5% carbon dioxide with the balance of nitrogen). The switch of the gas supplies was controlled by a 3-way valve with programming through uProcess scripting software (LabSmith., CA, USA). The interdigitated electrodes were soldered by copper-based wires on its soldering pads using the conductive epoxy adhesive (MG Chemicals, ON, CA) and connected to a commercial EIS HF2IS (Zurich Instruments, AG, Switzerland). Blood flow were observed using a CMOS camera at a frame rate of 10 frames per second (The Imaging Source, Charlotte, NC), which is attached to an Olympus X81 inverted microscope (Olympus America, PA, USA). Image contrast was enhanced by inserting a 414 ± 46 nm band pass filter in the optical path. The microfluidic device consists of a polydimethylsiloxane (PDMS) double-layer microchannel housing a microscale constriction matrix and interdigitated indium-tin-oxide (ITO) electrodes (Ossila Ltd, UK) (Fig. 1B and C). Fig. 1D shows the schematic of the core part of the microfluidic channel with capillary structures from our previous development.^4^ The interdigitated electrodes consist of 3 pairs of fingers with 100 µm band and 50 µm gap, well aligned to the intersectional area of the double-layer PDMS channels for optimal signal-to-noise ratio. Permanent covalent bonds were created between the PDMS channel and the glass substrate of electrodes using air plasma cleaner (Model PDC-001, Harrick Plasma) for 1 minute. Electrical impedance was measured using an AC signal at 2 Vpk at frequencies of 10 kHz, 50 kHz, 100 kHz and 500 kHz. Both components of the impedance, resistance *R*, and reactance *X*, were acquired at a sampling rate of 7 data points per second for analysis.

**Fig. 1.**
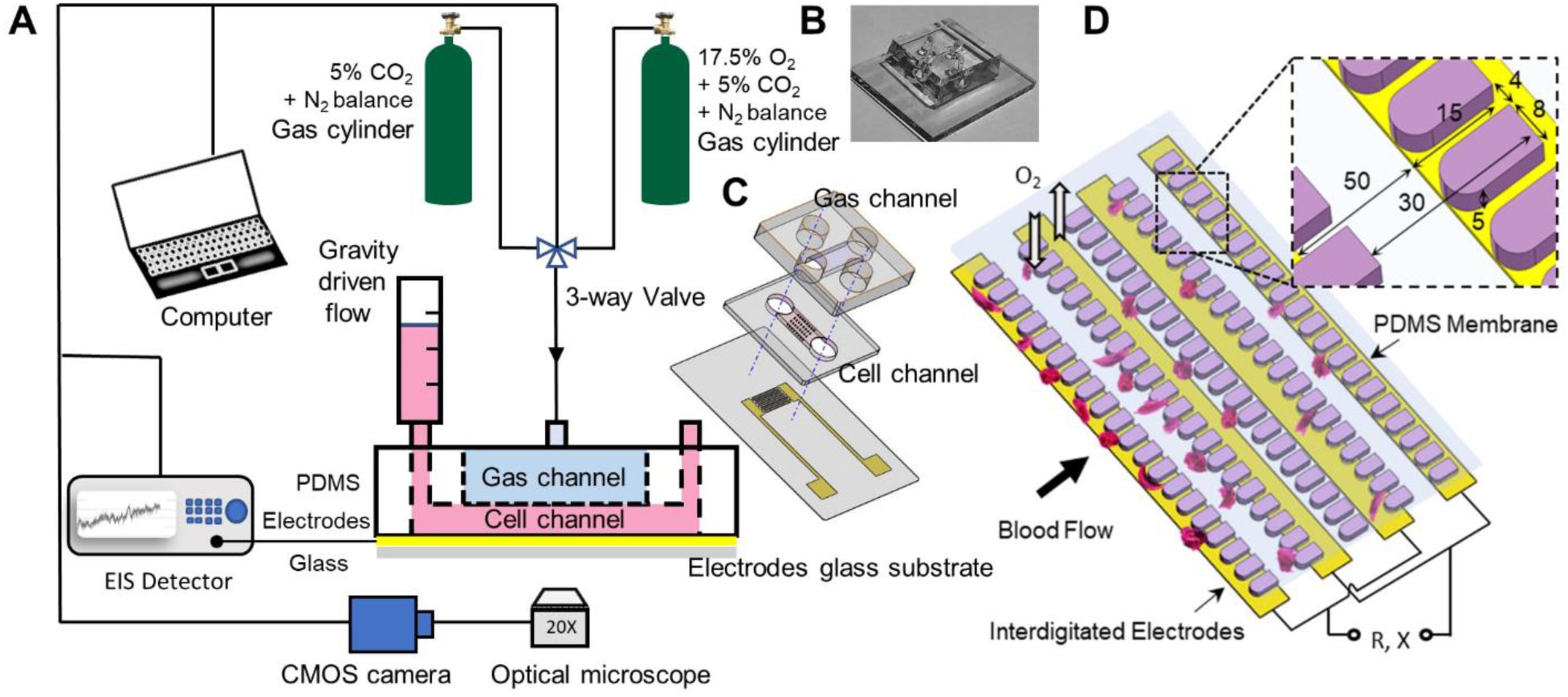
Microfluidic assay with electrical impedance spectroscope (EIS) detector for real-time vaso-occlusion testing. (A) A schematic of the complete experimental setup. (B) Exploded view drawing of the microfluidic device. (C) Schematic of the core part of microfluidic channel with capillary-inspired structures over the interdigitated electrodes. The *inset* shows the dimensions of the capillary structures.

## Results and discussion

Before the impedance measurement of vaso-occlusion, the experimental conditions of oxygenation in the gas channel and cell channel were calibrated using a fiber-optic O_2_ microsensor (Pyroscience GmbH, Aachen, Germany) connected to a FirestingO2 fiber-optic meter and controlled by Pyro Oxygen Logger software (Pyroscience GmbH, Aachen, Germany) (Fig 2. A). The microfluidic channels with equal experimental settings was flipped and bonded with and layer of PDMS on its top and glass slide on the bottom, which allowed the tip of the O_2_ microsensor to be easily inserted into the gas channel and cell channel (Fig 2. B). Fig 2. C demonstrates the results of O_2_ concentration profiles measured in the gas channel and cell channel during 3 cycles of switching of the gas supplies between high O_2_ concentration (17.5% O_2_ lasting 60 seconds) and low O_2_ concentration (0% O_2_ lasting 10 seconds). During the deoxygenation-oxygenation cycles, the O_2_ concentration varied between 17.1 % and 0.4 % in the gas channel, and the O_2_ concentration measured in the cell channel filled with PBS varied between 15.9 % and 0.5 %, respectively.

**Fig. 2.**
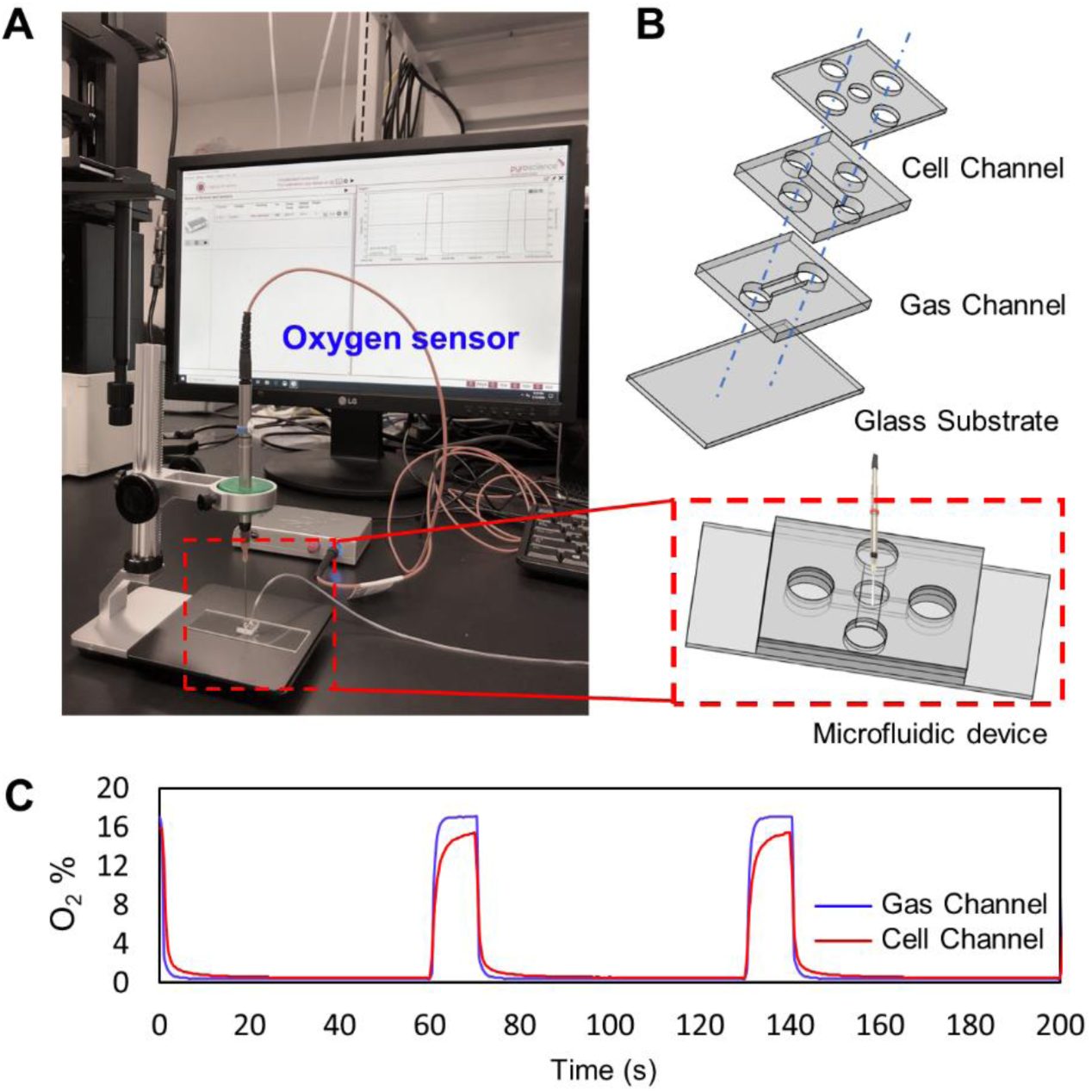
Oxygen concentration calibration within the microfluidic channels. (A) A photograph of the experimental setup of oxygen sensing in the microfluidic device. The *inset* shows the schematic of the microfluidic device (B) Exploded view drawing of the microfluidic device. (C) O_2_ concentration profiles measured in the gas channel and cell channel during cyclic switching of the gas supplies between high O_2_ concentration and low O_2_ concentration.

Performance of the developed vaso-occlusion assay was tested with normal (AA) and SS RBCs. As aforementioned, in order to further investigate the capability of drug efficacy testing of the assay, we next used the vaso-occlusion assay to examine the flow behavior of SS RBCs pretreated with anti-sickling agent of 5-HMF, which is known to have beneficial anti-sickling effect on SS RBCs by improving their oxygen affinity of hemoglobin.^18-20^ Impedance measurements and microscopic imaging process for validation were synchronously started upon the deoxygenation (lasting 60 seconds) immediately followed by reoxygenation (lasting 10 seconds) while the cells passing through the microvasculature-like microfluidic channel. Relative changes in both values of resistance (*R*) and reactance (*X*) were presented with their variations during the deoxygenation and reoxygenation processes (Δ*R* = *R* − *R*_0_, Δ*X* = *X* − *X*_0_). Impedance measurements were conducted at multiple frequencies simultaneously and repeated for each condition. Fig 3. A-D show the representative impedance signals measured at the frequency of 10 kHz. From the results, no distinctive change is observed in the impedance signals measured for AA RBCs during the deoxygenation and reoxygenation processes (blue curves). The corresponding time-lapse images demonstrate that deoxygenated AA RBCs were similarly deformable as in the oxygenated state, so that they were able to traverse capillary structures in the microfluidic channel regardless of the oxygenation condition (Fig 3. E and F). In contrast, drastic rise in the value of *R* and drop in the value of X were observed during the deoxygenation process of SS RBC flow; during the reoxygenation process, impedance signals reversed (orange curves).

**Fig. 3.**
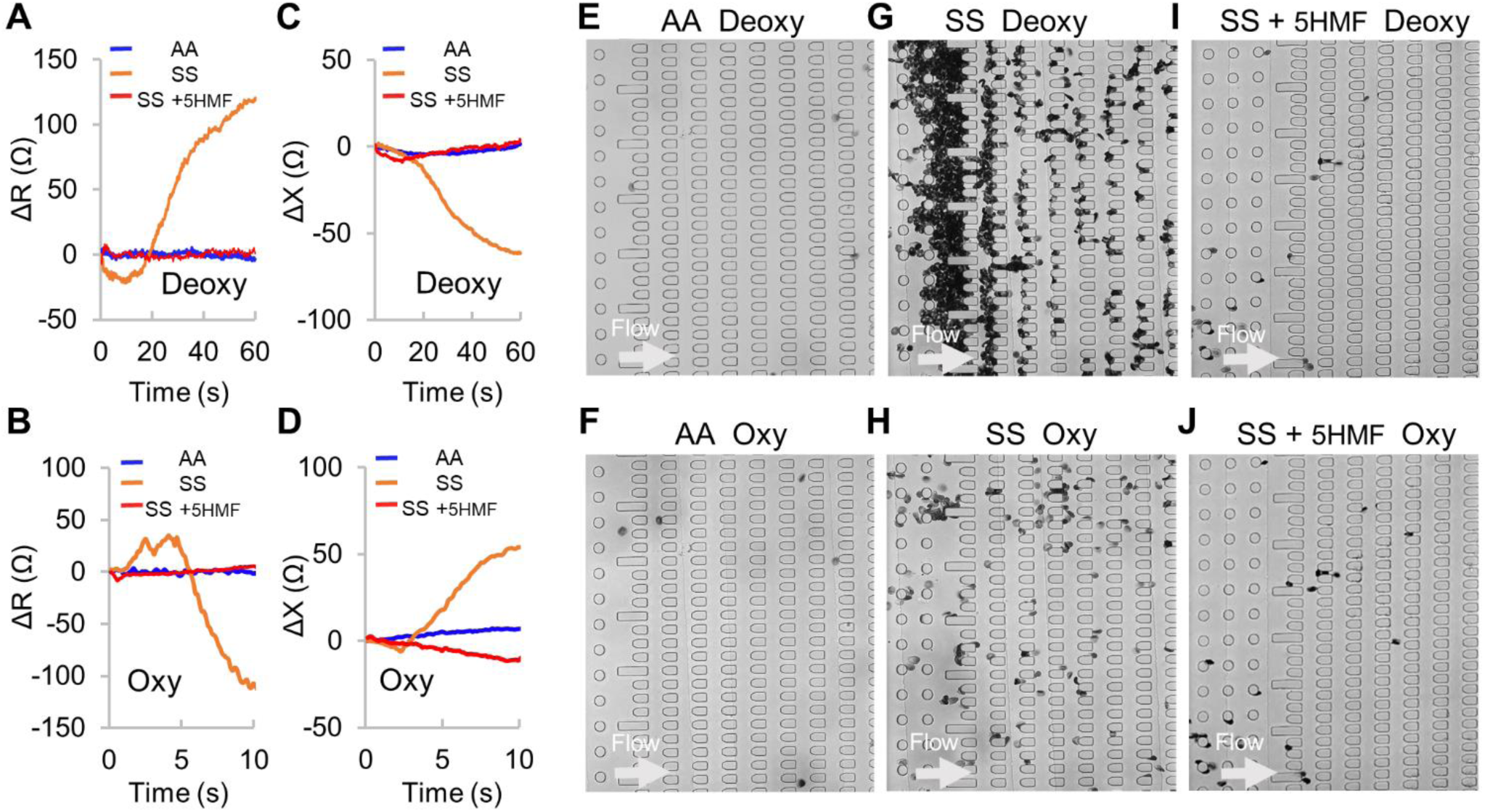
Real-time recordings of on-chip vaso-occlusion testing via electrical impedance and microscopic imaging. (A-D) Relative impedance signals measured at the frequency of 10KHz during the deoxygenation and reoxygenation processes. (E-J) Representative microscopic images of cells flowing through the micro-constrictions in the microfluidic channel. Normal RBCs – AA, sickle cells – SS, and sickle cells treated with 5-HMF– SS +5-HMF. The arrows indicate the flow direction.

Correspondingly, SS RBCs became sickled and trapped in the capillary structures during the deoxygenation process (Fig.3 G). SS RBCs experience intracellular HbS polymerization during the deoxygenation process, thereby making the cells more rigid.^4^ Such responsive reduction in deformability makes the cell more difficult to squeeze through the capillary structures, and ultimately leads to the cellular obstruction in the microfluidic channel.^4^ The resulting obstruction of deoxygenated SS RBCs consequently cause the shift of the overall electric impedance (increase of *R* and decrease of *X*) of the system. ^21^ When cells were reoxygenated, their deformability and the blood flow were subsequently resumed (Fig.3 H). In addition, we observed SS RBCs pretreated with 5-HMF shared similar behaviors to the AA RBCs in the rheology (Fig.3 I and J) as well as the resulting impedance signals (red curves), suggesting that the pretreatment with 5-HMF might greatly relieved the vaso-occlusion by sickle cells. These results demonstrate that the impedance measurement can reflect the cellular behavior in real time during the vaso-occlusion process in our assay. Fig.4 provides the complete data obtained at all the 4 frequencies under repeated deoxygenation and oxygenation conditions as indicated in Fig.2 C. The results show that the variations of the impedance signals follow similar trend among different cycles as well as different frequencies. As the frequency value increases, we can see the decrease in the sensitivity of the impedance measurement in response to the vaso-occlusion processes. Even though we didn’t find apparent obstruction of the blood flow during the cyclic deoxygenation-reoxygenation processes, there still existed variations of impedance signals for the AA RBCs and SS RBCs pretreated with 5-HMF, especially in the value of reactance at lower frequencies. This might be attributed to the influence of the noise from the gas pressure fluctuation due to the switch of gas supplies.

**Fig. 4.**
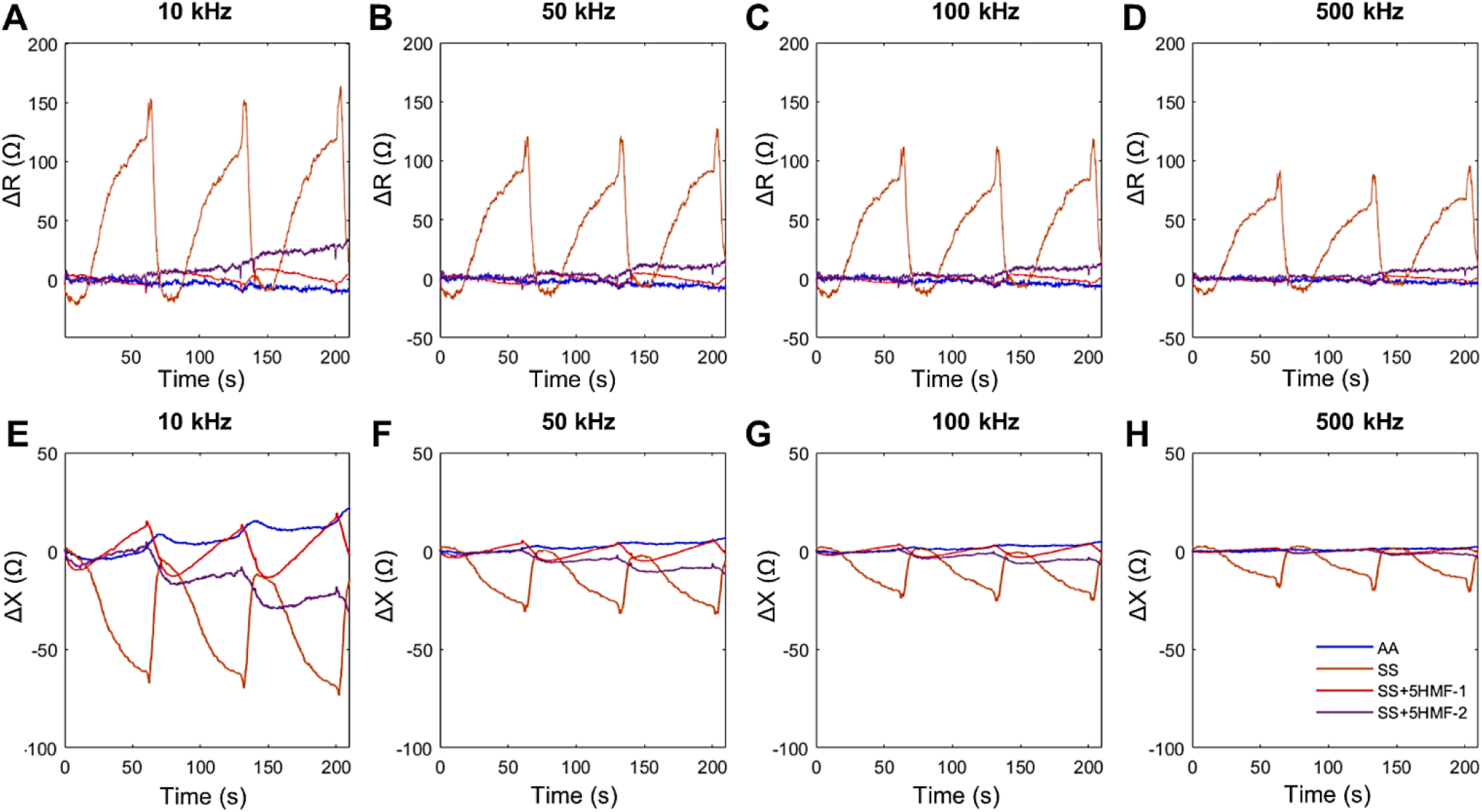
Relative impedance signals measured at multi-frequencies during the cyclic vaso-occlusion in the microfluidic channels. (A-D) Relative changes in the value of resistance (Δ*R*) measured at the frequencies: (A) 10KHz, (B) 50KHz, (C) 100KHz, (D) 500KHz. (E-J) Relative changes in the value of reactance (Δ*X*) measured at the frequencies: (E) 10KHz, (F) 50KHz, (G) 100KHz, (H) 500KHz. All the figures share the same legend. Normal RBCs – AA, sickle cells – SS, and sickle cells treated with 5-HMF– SS +5-HMF.

The unique feature of our assay is the integration of the electrical impedance sensing with the microfluidic mimics of capillary bed, which enables detection of blood flow and occlusion progress in real time. We envision this blood testing assay can be further miniaturized to a portable system for *in situ* assessment of vaso-occlusion progression and risk prediction of painful crisis from ultra-small volume of blood sample in sickle cell disease. The large benchtop equipment of the commercial impedance spectroscope can be substituted by a portable impedance device, enabled by an AD5933 impedance chip and an Arduino microcontroller, as used in a portable flow cytometry device.^17^ To be noted, our results have proven that we may obtain a more stable signal at a higher frequency during the impedance measurement in our experiment. Therefore, we may select the maximum operating frequency of AD5933 (100 kHz) as the optimal frequency of the portable system.^22^ The supply of hypoxia condition from gas tanks can be replaced by incubation of cells with oxidoreduction reagents, such as glucose oxidase (Sigma-Aldrich) and sodium hydrosulfite (Sigma-Aldrich).^12^ This developed assay can be potentially useful for monitoring blood flow conditions in SCD patients accepting pharmaceutical treatments, such as Hydroxyurea^23^ and L-glutamine^24^, as well as gene therapies.^25^

## Author contributions

Y.Q., J.L. and E.D. designed the experiments. Y.Q. and J.L. developed the device and performed the experiments. Y.Q. and J.L. analyzed the data. Y.Q., J.L., D.D. and E.D. discussed the results and constructed the manuscript.

## Acknowledgements

This work was supported by the NSF/IIS#1464102 and NIH/NHLBI OT2HL152638.

